# Long-read sequencing reveals extensive DNA methylations in human gut phagenome contributed by prevalently phage-encoded methyltransferases

**DOI:** 10.1101/2023.05.20.541561

**Authors:** Chuqing Sun, Jingchao Chen, Menglu Jin, Xueyang Zhao, Yun Y Li, Yanqi Dong, Na L Gao, Zhi Liu, Peer Bork, Xing-Ming Zhao, Wei-Hua Chen

## Abstract

DNA methylation is essential for the survival of bacteriophages (phages), yet, we know little about the methylation of their genomes. Here, we analyze the DNA methylation patterns of 8,848 metagenome-assembled high-quality phages across 104 fecal samples using single-molecule real-time (SMRT) sequencing. We show that 97.60% of gut phages could be methylated, and reveal factors that correlate with methylation densities. Phages having higher methylation densities are more prevent, suggesting putative viability advantages of the DNA methylation. Strikingly, more than a third of the phages encode their own DNA methyltransferases (MTases). More MTase copies are associated with increased genome methylation densities, methylation motifs, and higher prevalence of certain phage groups. Most MTases are closely homologous to gut bacterium-encoded ones, likely exchanged during phage-bacterium interactions, and could be used to accurately predict phage-host relationships. Taken together, our results suggest that the gut DNA phages universally use DNA methylation to escape from host defense systems with significant contribution from phage-encoded MTases.

## Introduction

Bacteriophages (phages) are viruses that infect bacteria and archaea[1], and modulate many ecological and evolutionary processes in complex microbial communities, including the human gut microbiome [2]. Phages often have narrow host ranges and are thus ideal tools for precision manipulation of the gut microbiota [3]. To defend the phages, prokaryotic organisms adopt a variety of defense mechanisms that are deployed against xenogeneic DNAs, among which the restriction-modification (RM) systems are ubiquitous and have been found in ∼90% of sequenced bacterial genomes [4]. RM systems often consist of a restriction endonuclease (REase) that recognizes a highly specific target DNA sequence (i.e., a distinctive, usually recurrent, molecular sequence, or motif) and degrades the unmethylated ones, and a corresponding DNA methyltransferase (MTase) that protects the same DNA sequence via the DNA methylation of the bacterial genome [4]. RM systems thus enable bacteria to distinguish self-genome from invading phage DNAs that are either unmethylated or not properly methylated. To escape from the host immunity, phages have developed strategies to overcome the RM systems, which can involve the methylation of their genomes through the hijacking of the host MTases, packing host MTase proteins into their virions, or incorporating host MTase genes into their genomes [5].

Recent exploration of large human virome/phageome datasets [6] have identified a substantial amount of novel gut phage genomes and revealed their diversity in human gut [2]. Furthermore, we have considerable anecdotal knowledge on the vital roles of phages in shaping the microbial community structure [5a, 7], mediating horizontal gene transfers among bacteria [8], and modulating host metabolic capacities [9]. In contrast, we know little about the epigenome landscape of entire phage microbial communities and the abundance of particular survival tactics, such as the encoding of their own MTases and the potential impact for the survival of phages.

Long-read sequencing techniques such as the single-molecule real-time (SMRT) and Nanopore have allowed us to explore the large-scale, genome-wide DNA methylations; they were successfully applied to genomes of eukaryotes [10] and prokaryotes [11], and recently also were used to characterize the methylation patterns in individual viral genomes [12] and those in environmental samples [13]. Although the Nanopore sequencing could identify more types of DNA methylations including the N^4^-methyl-cytosine (m4C), N^6^-methyl-adenine (m6A) and N^5^-methyl-cytosine (m5C) [14], the SMRT sequencing, especially in its circular consensus sequencing (CCS) mode could provide better resolution and higher accuracy (∼85%) in DNA methylation detection [15], and has been successfully used to characterize viral DNA methylations in marine samples [13a]. Hence, we applied SMRT sequencing to 104 viral-like particles (VLPs) enriched human fecal samples and conducted a comprehensive survey of the DNA methylation landscape of human gut DNA phages, which allowed us to identify and quantify DNA methylations and their contributing factors at a large scale.

## Results

### A set of high-quality phage genomes representing the human gut phageome

To obtain high-quality phage genomes for subsequent DNA methylation analysis, we first established a Chinese Human Gut Virome (CHGV) catalog consisting of 21,646 non-redundant phage genomes, via the combined assembly of short-(Illumina) and long-(PacBio in circular consensus sequencing (CCS) mode) reads (Methods; see also ref. [16]). Briefly, we enriched double-stranded DNA phages from fecal samples of 135 individuals of a Chinese population, subjected them to short-read Illumina sequencing, and selected 104 samples with sufficient amounts of high-molecular weight DNA for PacBio SMRT long-read sequencing (Methods). We reconstructed the phage genomes through a hybrid assembly pipeline using both the short- and long-reads, followed by de-replication, and viral genome recognition to generate a non-redundant set of phage genomes (Fig. 1A) [16]. The viral recognition procedure included viral prediction by VirSorter [17], VirFinder [18], PPR-Meta [19], protein annotation against the NCBI POG (Phage Orthologous Groups) database [20] and comparing the assembled contigs to NCBI Viral RefSeq using BLASTn [21], followed by evaluation of the outputs based on six commonly used criteria (Methods).

**Figure 1.**
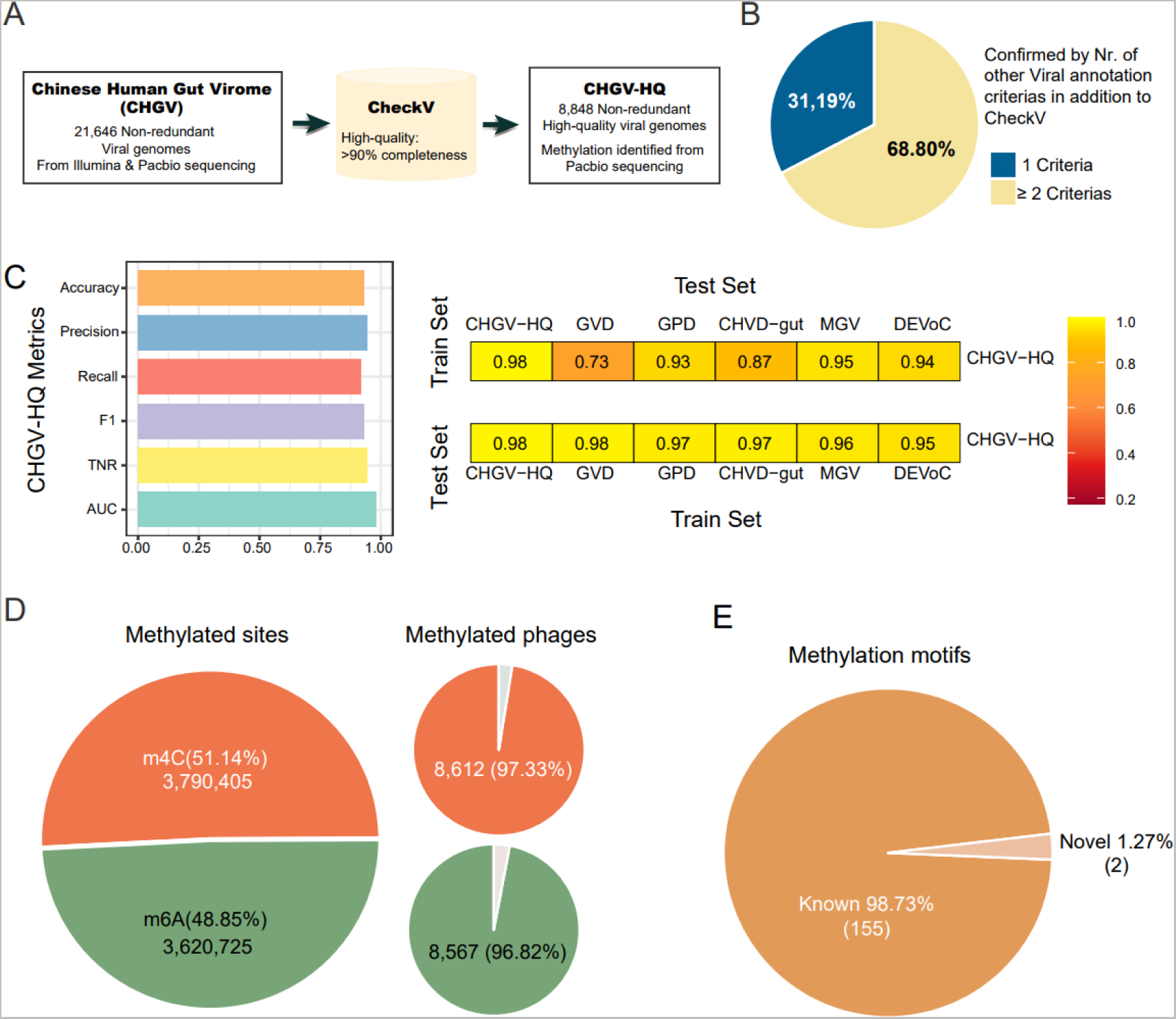
The DNA methylation landscape of the human gut phageome identified using a representative set of 8,848 high-quality gut phages. **A**) Generation of 8,848 high-quality non-redundant double-stranded DNA phages from human fecal samples (CHGV-HQ) using combined assembly of short-(Illumina) and long-(PacBio)read sequencing. **B)** Percentages of CHGV-HQ genomes annotated by the numbers of viral-recognition criteria in addition to CheckV (Methods). **C**) Left panel: Performance metrics of the viral-prediction machine learning model based on the CHGV-HQ genomes (i.e., the CHGV-HQ model) in distinguishing the viral genomes in the IMG/VR database from the bacterial genomes in the UHGG2 database (Methods). Y-axis: Accuracy – correct predictions out of all predictions, Precision – ratio of true positives over the sum of false positives and true negatives, Recall – correctly predicted outcomes to all predictions, F1 – combines the accuracy, precision, and recall metrics into one single metric that ranges from 0 to 1, TNR – true negative rate, AUC – **a**rea **u**nder the ROC (receiver operating characteristic) **c**urve. Right panel: Proportions of correctly recognized phage sequences in CHGV-HQ and public virome databases by the viral-cognition machine learning models. Top: proportions of the phage genomes in public virome databases that were correctly recognized as phages by the CHGV-HQ model. Bottom: proportions of the CHGV-HQ phage genomes that were correctly recognized by ML models based on the public virome databases (Methods). **D**) DNA methylation sites and their prevalence in the 8,848 phages, stratified by methylation types, including N6-methyl-adenine (m6A or 6mA) and N4-methyl-cytosine (m4C or 4mC). **E**) Methylation motifs identified in this study and their overlaps with those in the REBASE (http://rebase.neb.com/rebase/rebase.html).

To avoid biases brought by the fragmented phage genomes, we selected a subset of 8,848 high-quality phages with ≥90% completeness (CHGV-HQ hereafter; Table S1) according to CheckV[22]; in addition to the CheckV recognition, 68.80% of the genomes were recognized by at least two viral recognition criteria (Fig. 1B).

We next examined whether the CHGV-HQ genomes could allow us to obtain an unbiased view on the DNA methylation landscape of the human gut phageome. Because the human gut phageome was both diverse and individual specific [2], rather than directly comparing the phage sequence to those in public human virome databases, we instead checked if our CHGV-HQ phage genomes represented most of the sequence signatures of known human gut phages. Briefly, we trained a virus detection machine learning (ML) model using the 8,848 genomes as the true positives and a subset of the Unified Human Gastrointestinal Genome (UHGG) [23] genomes as true negatives (the CHGV-HQ model; Methods). We first tested the model on an independent dataset consisting of viral genomes from the IMG/VR database [24] and the left-out bacterial sequences in the UHGG. The model could accurately distinguish the viral from bacterial sequences with an overall area under the receiver operating characteristic curve (AUC) value of 93% (94% precision and 92% recall rates; Fig. 1C). We then applied the model to the public virome databases including the Gut Virome Database (GVD) [6a], Gut Phage Database (GPD) [6b], Metagenomic Gut Virus catalog (MGV) [6c], Cenote Human Virome Database (CHVD) [6d], and Danish Enteric Virome Catalog (DEVoC) [6e]. The CHGV-HQ model correctly recognized ∼88% (73%∼95%) of the viral genomes at a false-discovery rate of 5.72%, suggesting that the CHGV-HQ catalog captured the key sequence characteristics of human gut virome/phageome. In addition, virus-detection ML models trained on the true positives from the public virome datasets correctly recognized ∼96% (95%∼97%) of our CHGV-HQ genomes as viral genomes (Fig. 1C), further confirming their viral identity and demonstrating that the sequence signatures were sufficient for further analysis.

### The DNA methylation landscape of the human gut DNA phageome

We analyzed the long-read sequencing data in the 104 samples (Fig. 1A; Methods) and identified DNA methylation sites using the SMRTlink tool, which had a ∼85% detection rate and ∼95% accuracy according to its manual [15]. For each sample, we aligned the consensus reads (HIFI reads) of the subreads generated by the PacBio CCS mode to the 8,848 CHGV-HQ phage genomes and then used the SMRTlink tool to detect the methylated bases which compared the mean Inter Pulse Duration ratio (IPDr) of all the subreads at a position of the reference genome with that of the unmethylated bases [15]. We obtained a total of 8,448,009 non-redundant DNA methylation sites, among which 50.73% and 49.27% were m4C and m6A modifications respectively. In total, 8,630 (97.53%) out of the 8,848 phages were methylated, and 8,549 (96.6%) contained both types of modifications (Fig. 1D).

We identified a total of 157 DNA methylation motifs using an established method MEME tool [25] based on the large number of phages and identified methylation sites. Among which, 115 and 100 were m4C and m6A motifs, respectively, 58 motifs were shared by both methylation types, i.e., same motif sequence with different methylated base (Table S1; Methods). Of which, 155 motifs were identical to the REBASE motifs [26] (Fig. 1E; Methods).

We then characterized the methylation patterns of the phage genomes, and identified genomic and evolutionary features that favored them. To better capture signatures of the DNA methylation in different genomes, we combined the methylation sites of all the CHGV-HQ genomes from all samples (Fig. S1), assuming that all potential methylation sites could be methylated in a genome. We observed a lower density of the m4C methylations in the coding regions (as measured by the proportion of cytosines methylated per genome) than the non-coding regions, but an opposite pattern for the m6A methylations (Fig. 2A, Table S1; p < 0.001, Wilcoxon Rank Sum Test; Methods). We further dissected these patterns by assigning the phages to four lifestyle groups, namely temperate, uncertain temperate, uncertain virulent and virulent using a DeePhage tool [27], and observed an increasing trend in the m4C densities with the increasing phage virulence, i.e., increased from temperate to virulent in both coding and non-coding regions, but an opposite trend in the m6A methylation densities (Fig. 2A). Overall, we found significantly higher methylation density of m4C than m6A in both coding and non-coding regions. However, further analysis revealed that the differences in the coding regions were caused by lower proportions of m6A-methylated coding genes (CDSs with at least one modified A base, Fig. 2B; *p* < 0.001, Wilcoxon rank sum test) rather than lower m6A densities in each of the CDSs (Fig. 2C; *p* >0.05, Wilcoxon rank sum test). Additionally, we also observed a correlation between the phage lifestyle with the proportion of methylated CDSs. In particular, the proportion of m4C methylated CDSs increases with increasing phage virulence, while the opposite pattern was found for m6A (Fig. 2B). These results indicate different methylation pattern preference in gut phages between lifestyle groups. Previous studies have shown that the phage lifestyle is a determinant factor shaping the phage defense strategies against the bacterial hosts [28], our results thus suggested the two methylation types (i.e., the m4C and m6A methylations) may play different roles in phages of different styles.

**Figure 2.**
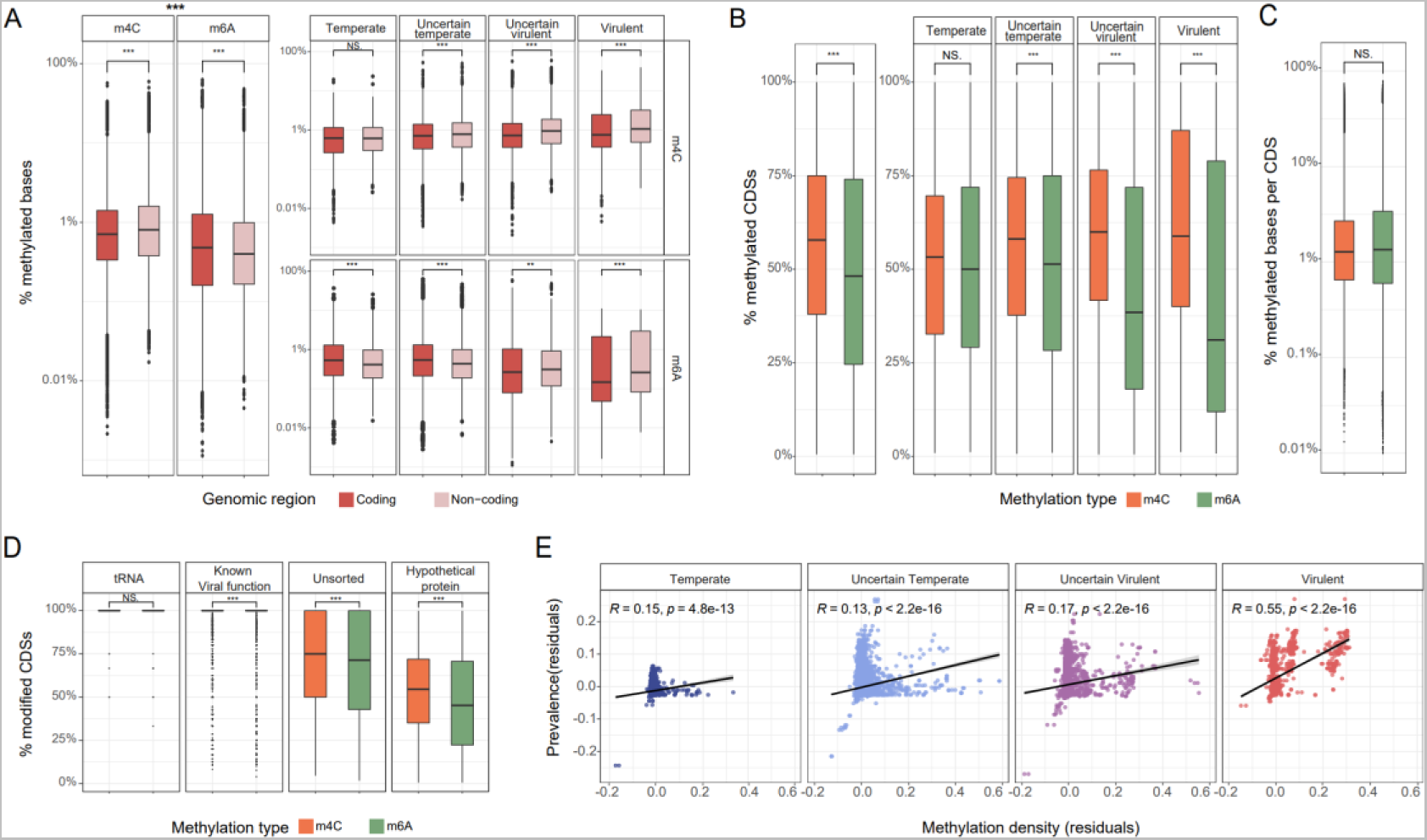
Distributions patterns of the DNA methylations in CHGV-HQ genomes as functions of phage lifestyles, gene functions and phage prevalence. **A**) Methylation density with different distribution patterns of m6A and m4C in coding versus noncoding regions, and the impacts of phage lifestyles. The methylation density was calculated by number of methylated bases divided by total number of the related bases, e.g., number of mA bases/number of A bases. The overall trend of coding region being enriched in m4C methylation remained in each lifestyle group (right-top panel). Conversely, the m6A densities decreased with the increasing phage virulence (right-bottom panel); however, because the m6A densities decreased further in the coding than the noncoding regions in the two virulent phage groups (i.e., uncertain virulent and virulent), we found significantly higher m6A densities in the coding than the noncoding regions of the two virulent groups (right-bottom panel). We found significantly higher m4C densities in both coding and noncoding regions than that of m6A (left panel). **B**) Differential distribution patterns between m6A and m4C in coding sequences (CDS), and the impacts of phage lifestyles. The number methylated CDSs takes all the CDSs with at least one correlated methylation site into account. **C**) The ratio of modified bases/genome length per CDS between m6A and m4C. **D**) Differential distribution patterns of m6A and m4C modifications in coding genes with different functions; almost all tRNA genes and coding genes with known viral functions (Including Assembly, Immune evasion, Lysis, Integration, Replication, Regulation, Packaging, and Infection) are methylated, as compared with much lower methylation rates in phage genes coding for unsorted and hypothetical proteins. See Fig. S2 for more detailed gene functional categories. **E)** The viability of the CHGV-HQ phages, i.e., the prevalence across 104 fecal samples, was positively correlated (Partial correlation using Pearson correlation, *p* < 0.001) with the overall DNA methylation density when the sequencing depth was controlled for (Methods). Prevalence was calculated by using an abundance cutoff of 0.5 RPKM as the presence/absence threshold (Methods); the results obtained when using other abundance cut-offs as the threshold were similar and can be found in Fig. S3E, F. Plotted here are the residuals of the prevalence (Y-axis) and methylation density (X-axis) after the sequence depth was controlled for (Methods).

We also observed differential methylation patterns in genes with various functions (Fig. 2D). For example, tRNA genes, and most genes encoding proteins with known viral functions such as roles in lysis, immune evasion, integration, assembly, packaging and infection, were methylated by both m4C and m6A in most genomes, in contrast to significantly lower proportions observed for genes with less characterized functions such as those coding for hypothetical and unsorted proteins (Fig. 2D and Fig. S2). These results suggest that functionally important genes are more likely to be methylated.

We also examined whether the methylation densities could correlate with the viability of the gut phages. The latter could be evaluated from two aspects, the ability of a phage to accumulate in one sample (i.e., abundance), and the ability to spread and survive across samples (i.e. prevalence). Because the methylation density is significantly affected by the sequencing depth (e.g., the accumulative coverage of the CHGV-HQ genomes in all samples) and the abundances (i.e., the higher the abundance, the higher the coverage), we thus calculated a partial correlation between the phage prevalence and the methylation densities, while controlling for the sequencing depth. We determined the prevalence of the 8,848 CHGV-HQ phages in the 104 samples using an arbitrary relative abundance cutoff of 0.5 calculated from the short-read viral-like particles (VLP) sequencing data (Methods), and observed a strong positive correlation between the prevalence and methylation densities (Fig. S3A; *p* < 0.001, Partial Pearson correlation *r* = 0.35 after the sequencing depth was controlled) as well as the methylation motifs (*p* < 0.001; Fig. S3B). Because virulent phages are generally more abundant and prevalent than the temperate phages [6a], we also stratified our analysis according to the phage lifestyle groups and found similar trends (Fig. 2E, Partial Pearson correlation after the sequencing depth was controlled). The same trends were found for both the m4C and m6A methylation types (Fig. S3C, D). In addition, changing the presence/absence threshold to other arbitrary abundance cut-off in our prevalence calculation did not affect our main results (Fig. S3E, F). To further remove discovery biases, we also limited our analysis to phages with more than 100× coverages, and obtained similar results (Fig. S4A, B). Thus, our results suggested that higher methylation densities and/or numbers of methylation motifs might correspond to higher phage viability.

Together, our results show that the DNA methylation is a universally present in gut phages and may play important roles for their survival in the human gut.

### Phage-encoded MTases are prevalent and associated with higher DNA methylation density and phage viability

To clarify the mechanisms underlying these frequent methylations, we examined whether gut phages encode their own MTases. We searched all annotated proteins of the 8,848 CHGV-HQ phage genomes against the Conserved Domain Database (CDD) [29] using RPS-BLAST[30] (v2.12.0+) (Methods), and identified a total of 4,064 putative MTases with an E-value cut-off of 1E-5. Overall, phage-encoded MTases are prevalent, and could be found in 34.09% (3,205, Fig. 3A) of the CHGV-HQ phages (Table S1). Among which, 72% of the MTase-containing phages encoded only one MTase gene, whereas the others harbored multiple such genes (Fig. S5).

**Figure 3.**
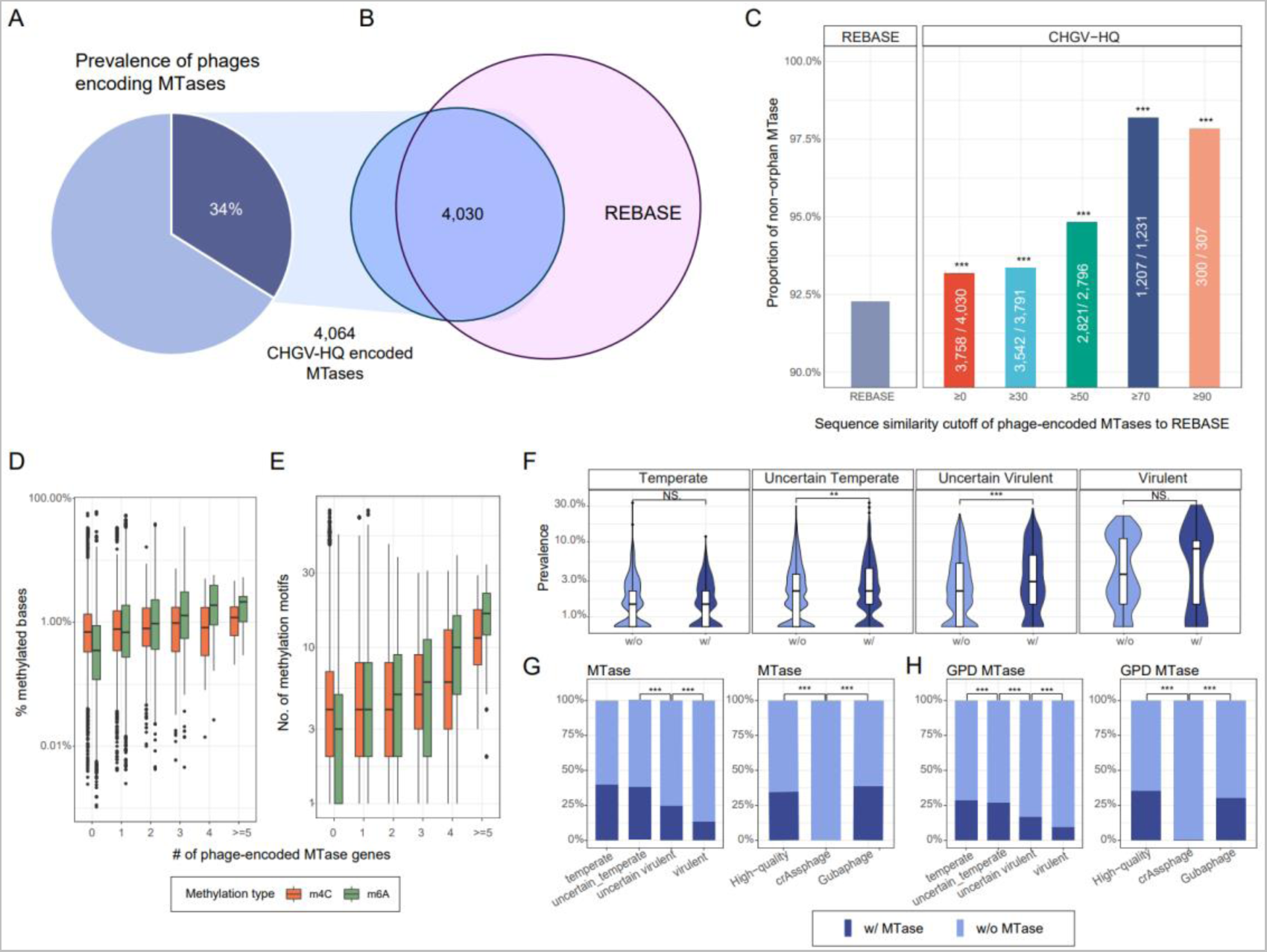
Phage-encoded MTases are prevalent in gut phages, and are associated with higher phage viability. **A)** Over one-third of the CHGV-HQ phages encode their own MTase genes. **B)** Most (4,030 out of 4,064, 98%) of the phage-encoded MTases share significant protein similarities with those in the REBASE (BLASTp e-value <1e-5; Methods). ≥0 stands for all phage-encoded MTases with BLASTp e-value <1e-5. **C)** The phage-encoded MTases are mostly non-orphan MTases, and contain significantly higher proportion of non-orphan MTases than the REBASE. Chi-square tests were performed between REBASE and each of the CHGV-HQ groups with different protein similarity cutoffs; \**p* < 0.05, \*\**p* < 0.01, \*\*\**p* < 0.001. **D, E)** Increasing numbers of phage-encoded MTase genes are associated with higher methylation densities and more methylation motifs in the phage genomes. **F)** Violin plots showing prevalence of the phages with/without MTases (X-axis), stratified by the phage lifestyles (i.e., from temperate, uncertain temperate, uncertain virulent, and virulent). Wilcoxon rank sum tests were performed between groups; \**p* < 0.05, \*\**p* < 0.01, \*\*\**p* < 0.001. **G)** Bar plots on the left showing the prevalence of MTase genes in the CHGV-HQ phages as a function of phage virulence: a decreasing MTase prevalence is obvious with the increasing phage virulence (i.e., the highest in temperate phages, and lowest in virulent phages). Similar trends could be found in individual MTase types (Fig. S6B, C). Bar plots on the right showing the heterogeneous distributions of the MTases in phage subgroups such as the crAssphages and Gubaphages. The same trends could be found in GPD phages **H)**.

In addition to the RM systems, DNA MTases may exist without cognate REases, in which case they are referred to as the orphan MTases, and likely involved in processes other than anti-phage immunity such as gene expression regulation [31], and DNA replication and repair [32]. We thus searched the phage-encoded MTases against those in the REBASE database that were already classified as orphan or non-orphan MTases. Out of the total 4,064 MTases, 4,030 had significant BLASTp hits with an E-value cut-off of 1E-5; among which, 93.25% (3,758) had BLASTp hits with the non-orphan MTases, significantly higher than the overall 92.27% non-orphan MTases in the REBASE (207,303 out of 224,651; Fig. 3C, *p* < 0.001, Hypergeometric test). Furthermore, the phage-encoded MTases with higher protein similarities (e.g., ≥90%) with those in the REBASE were further enriched with the non-orphan ones (97.71%, Fig. 3C, *p* < 0.001, Hypergeometric test). Thus, most of the phages-encoded MTases are likely to be involved in the phage-bacterium defense processes.

We observed an increasing methylation density and number of motifs in the phages with more MTases genes (Fig. 3D, E; see also Fig. S6A for the trends for the individual modification types), suggesting that these phage-encoded MTases were indeed functional and contribute to the methylation of their encoding phages. In addition, we also observed that the phage-encoded MTases contributed to significantly higher prevalence in two out of the four lifestyle groups (Fig. 3F; the prevalence was measured by using an arbitrary abundance cut-off of an RPKM≥ 0. 5 as the presence/absence threshold across the 104 samples; Methods). These results suggest that the phage-encoded MTases may also contribute the spreading of the corresponding phages across humans.

Phage lifestyle was also associated with the distribution of the MTase genes. For example, we found a significantly higher prevalence of MTase genes in temperate phages than in the virulent ones (Fig. 3G, H). The trends in the individual MTase types, i.e., MTases responsible for m4C and m6A modifications were mostly the same (Fig. S6B, C, line charts; Table S2). The higher occurrence of MTases in temperate phages is likely due to their increased time spend within host cells and hence higher chances in exchanging genetic material horizontally.

However, heterogeneity in the lifestyle distributions was observed among different taxonomic groups. For example, although both crAssphages and Gubaphages, the two most prevalent viral clades in human gut, are known to be virulent[6b], none of the crAssphage genomes in our CHGV-HQ collection encode MTase genes as compared to ∼38.9% of the Gubaphages (*p* < 0.001, Chi-squared test). Similar trends were found among the high-quality gut phage genomes in the Gut Phage Database (GPD [6b]; Fig. 3H; Methods), implying that crAssphages developed other means of achieving high prevalence besides avoiding the RM-systems.

In summary, our results show that over one-third of the CHGV-HQ phages encode their own MTase genes, which are associated with higher DNA methylation density and increased phage viability.

### Phage MTases are closely homologous to bacterium-encoded ones and can be used for accurate phage-host prediction

To further characterize the phage encoded MTases, and to quantify previously observed similarity with bacterial genes [5, 33], we annotated MTases from the Unified Human Gastrointestinal Genome v2.0 (UHGG2) [23], a metagenome-assembled human gut microbial genome collection by applying the same pipelines used for CHGV-HQ (Methods). We then combined the 61,719 annotated MTases with the 4,064 phage-encoded ones identified in the CHGV-HQ genomes, performed all-against-all protein similarity searches using BLASTp[30], and built protein clusters (PCs) using a Markov clustering algorithm (MCL) [34]. We obtained a total of 1,409 MTase PCs (including singletons), among which 7, 1,228 and 174 clusters contained phage, bacterial, and both bacterial and phage MTase genes; we referred to the last group of PCs as gene-sharing PCs (GS-PCs). Importantly, 4,048 (99.61%) of the total phage MTases (v-MTases) were included in the GS-PCs, (Fig. 4A) among which, 59.38% of the v-MTases share over 90% protein sequence identities with their bacterium-encoded homologues (Fig. 4B). We observed similar results between phage- and bacterial-encoded MTases using experimentally validated phage-host relationship data from the Microbe-versus-Phage (MVP)[3] database (Fig. S7A). The high similarity of the gut phage MTases with bacterium-encoded ones thus points to not only a frequent gene exchange but also potential bacterial hosts.

**Figure 4.**
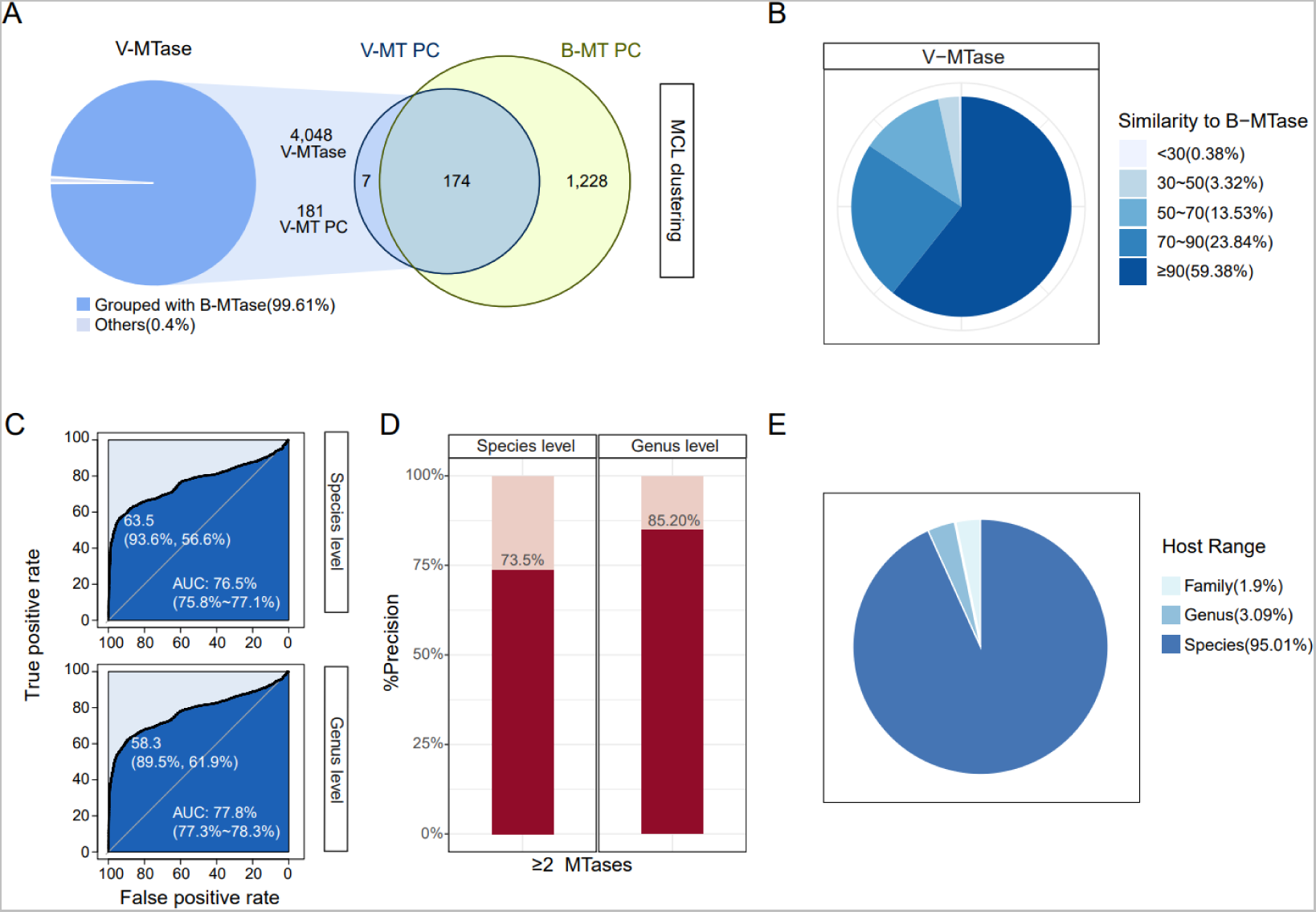
Phage-encoded MTases are homologous to bacteria-encoded ones, and can be used to predict phage-host relationships with high precision. **A)** Most phage MTases (v-MTases) can be clustered into protein clusters (PCs) together with their bacterial homologs (b-MTase) using a Markov Cluster Algorithm (MCL). **B)** Distributions of protein sequence similarities of phage MTases. **C)** Phage-host prediction AUROC in the golden-standard dataset using at least one MTase at species- (Upper) and genus- (Lower) levels. **D)** Phage-host prediction precision in the golden-standard dataset using the two or more MTase at species- (left) and genus- (right) levels. **E)** Host ranges for phages with predicted hosts using MTase.

To validate the exchange of MTases indeed occurred between phages and their hosts, we examined whether such genes could be used to accurately distinguish phage-host relationships in the MVP database from randomly generated phage-bacterium pairs (Methods). By using the BLAST results between phage and bacterial MTases as the only feature, we achieved overall accuracies of 76.5% and 77.8% area under the receiver operating characteristic (AUC) values at the species and genus levels, respectively (Fig. 4C). Using at least one MTase, the phage-host relationships could be predicted at 67.20% and 82.08% precision at the species and genus levels, respectively. The prediction precision could be further increased to 73.5% and 85.2% (Fig. 4D), by requiring that at least two MTase genes were shared between the phage-bacterium pairs with protein similarity higher than 90% (Methods). At the latter criteria, we could predict a total of 4.77% phages to their hosts (Table S1). We further validated the prediction results by calculating the host range for each of the phages as the last common ancestor (LCA) of all predicted bacterial hosts passing the above criteria, and found that 95.01% of the phages whose hosts had LCA at the species level (Fig. 4E, Fig. S7B). Together, these results support the assumption of the frequent exchange of MTases between phages and their hosts. Since the exchanges are bidirectional [35], further work is needed to quantify directionality, as acquisition by phages increase their viability, the transfer of MTases to host bacteria could also confer changes through the increased DNA methylation potential.

## Conclusion

In this study, we use the SMRT technology to characterize the first DNA methylation landscape of human gut phages at the community scale. Our finding that the gut phages may use DNA methylation to escape bacterial anti-viral immune mechanisms has several important implications. First, the similar analysis could be applied to the phage communities in other environments [13a] to better illustrate the role of methylation and phage-encoded MTases in those environments. Since the RM systems are universally present in bacterial genomes, survived phages appear to encode the respective counter-mechanisms, among them the DNA methylation. Second, the finding that differential methylation patterns exist in genes with various functions suggests that for some genes, by shielding themselves with DNA methylation, are able to keep functioning in the bacterial genome after the phage infection. Furthermore, the higher methylation densities found in more virulent phages indicate that DNA methylation is crucial for phages to escape the restriction endonuclease recognization from their hosts. It has been suggested that the phage-encoded MTases, which are mostly orphan ones, can be functional methyltransfereases and help the phages to overcom the bacterial R-M Systems and even contribute to the emergence of phages with broad host ranges [36]. The fact that these genes are different of their methylation pattern further indicate that they are of important functions, such as those with known viral functions including the infection, immune evasion and assembly. Third, the phage-encoded MTase shared significant protein similarity (often >90%) to bacterial encoded ones, indicating that they were frequently exchanged between phages and bacteria and thus could be used to infer phage-host relationships. Sequence similarity-based approaches have been used to establish such relationships in many environments [37]; consistently, our analysis showed high precision in predicting phage-host relationships using MTases in an experimentally validated dataset (Fig. 4D). Last, the viability benefits that we observed in phages with self-encoded MTases might represent only a small part of the functional consequences of the large numbers of genes frequently exchanged between phage and their bacterial host genomes. We found that there are ∼20% genes share high similarity with bacterial genes (Fig. S8), suggesting that there might be other genes that help the virus survive in the bacteria, and these genes could also be used to predict the host.

Our estimation on the extent of the phage DNA methylation and annotation of MTases may suffer from a few technical drawbacks. First, although the PacBio CCS mode allowed us to more reliably detect methylation signals from the subreads, i.e., a genomic fragment from a single virion was sequenced multiple times and thus the consensus signals were more reliable than using reads from multiple virions of the same species/strains, heterogeneous DNA methylation patterns due to the high micro-diversity caused by high viral mutation rates and/or partial methylation within a viral population could lead to increased false negative calls. The distribution of methylation density further confirmed that very few methylation-positive genomes with low methylation density (Fig. S9), thus reduces the possibilities of the overestimate methylation prevalence due to the misalignment of the identified methylation fragment. In addition, phages infecting multiple bacterial species may have different methylation patterns from different hosts, although such cases could be rare because most phages have rather narrow host ranges at the species and even strain levels [3]. Sequence depth is critical factor in methylation identification, too. Our rarefaction analysis indicated the number of unique methylation density increased with the increasing sequence depth and the rarefaction curve is far from saturation (Fig. S4A). Thus, our estimation may present only part of the global picture of the full extent of the phage DNA methylation landscape in the human gut. There is still in need of a more reliable bioinformatics pipeline to identify methylation sites in genomes outside the eukaryotic and prokaryotic worlds. Second, our annotation of MTases was rather conservative. In this study, we used an RPS-BLAST based search that is known to less sensitive than hidden Markov (HMM)-based search methods [38]. However, the latter might suffer from high false positives [39]. Last, we identified too many sequence motifs in the gut phages. For example, although we obtained on average 4∼5 motifs per phage, some phages could contain more than 30 motifs. This was in part because that some MTases are not site specific [40], or the less stringent search criteria used by MEME.

Interestingly, we observed a significantly different distribution of the two modification types, namely m4C and m6A between the gut phageome and bacterial genomes. For example, the two methylation types accounted for 50.75% and 49.25% in our dataset, which was in sharp contrast to the bacteriome that the m6A is dominant and accounts for ∼75% of the modifications [41], whereas m4C accounts for ∼20% [42]. The discrepancy could be due to biological reasons such as that some bacteria are associated with more phages than others thus causing the overall methylation patterns in phages to biased towards those bacteria, or technique reasons such as detection biases because of the lack of dedicated viral methylation identification tools. Of note, we did not analyze m5C modifications because of two reasons. First, currently the SMRT sequencing could not reliably detect them for prokaryotes and phages [15]. Second, m5C modifications are known to be rare in prokaryotic genomes [42]. Regardless, our knowledge on methylation patterns of the human gut bacteriome so far came from only a few studies with few fecal samples [41] and a comprehensive understanding is still lacking. In summary, we systematically characterized the DNA methylation landscape of human gut phages at the community scale using PacBio SMRT sequencing on 104 VLP enriched samples. Our results suggest that the gut DNA phages universally use DNA methylation to escape from host defense systems with significant contribution from phage-encoded MTases. Together with recent studies exploring the identification of human phages at large scales [6a-e], our data and findings have only started to picture the survival tactics of the gut phages, the functional consequences of the frequent phage-bacterium gene exchanges to shape the gut microbial communities, and more daring, their implications in human health and diseases [43].

## Methods

### Generation of 8,848 near-complete double-stranded DNA phages from the human gut

The Chinese Human Gut Virome (CHGV) catalog was obtained from a previous study [44] that contains 21,646 non-redundant viral contigs generated by combined assembly of short- (Illumina) and long-(PacBio) reads. Briefly, a total of ∼500g of feces from each of the 135 healthy volunteers of Chinese residence were collected and processed by a virome enrichment protocol to obtain a large number of viral-like particles (VLPs). High-quality and high-molecular weight double-stranded DNAs were then extracted, and submitted to Illumina HiSeq2000 sequencer (Novogen, Beijing, China) for viral next generation sequencing (vNGS, short-reads). A subset of 104 samples with sufficient quantity of viral DNAs was also submitted to the PacBio RS II sequencer (Pacific Biosciences, Menlo Park, CA, USA) with Circular Consensus Sequencing (CCS) mode for viral third-generation sequencing (vTGS, long-reads). Human genome contaminations were identified and removed from both the vNGS and vTGS datasets, followed by a combined assembly pipeline to generate putative phage contigs (see ref. [44] for more details).

To identify viral genomes, the following tools were used: VirSorter v2.0 [17] (--min-score 0.7), VirFinder v1.1 [18] (default parameters), and PPR-Meta v1.1 [19] (default parameters). A BLAST search against the Viral RefSeq genomes was also performed using BLASTn v.2.7.1 [21] with the default parameters and an *E*-value cutoff of <1e-10; Release 201 (Jul 06, 2020) of the Viral RefSeq database contained 13,148 viral genomes. In addition, the annotated protein sequences were used for BLAST searches against the NCBI POG (Phage Orthologous Groups) database [20].

A contig was annotated as a virus if it was circular/met at least two of the following criteria 1-5, the same criteria have been adopted by the GVD [45]:

1. VirSorter score ≥ 0.7,
2. VirFinder score > 0.6,
3. PPR-Meta phage score > 0.7,
4. Hits to Viral RefSeq with > 50% identity & > 90% coverage,
5. Minimum of three ORFs, producing BLAST hits to the NCBI POG database 2013 with an *E*-value of ≤ 1e-5, with at least two per 10 kb of contig length.
6. Alternatively, contigs met one of the above criterium and were annotated as high-quality (≥ 90% completeness) by CheckV [46] were also annotated as viruses.

To avoid bacterial contamination, we first identified possible prophage regions using PhageFinder [47] (v2.1) and removed them from UHGG genomes to prevent over-estimation of the contamination. The resulting UHGG dataset was referred to as UHGG-Minus in this study. We then carried out a BLAST search against the UHGG-Minus sequences using BLASTn v.2.7.1[21] with the default parameters and an *E*-value cutoff of <1e-10, and contigs with blastn hit of 90% identity over 50% of its length were removed from further analysis. To avoid fragment genomes, the contigs were filtered with length longer than 5 kb or circular contigs longer than 1.5 kb.

In the end, a non-redundant set of 21,646 viral contigs was obtained and referred to as the Chinese Human Gut Virome (CHGV) catalog.

The 8,848 near-complete double-stranded DNA phages were then screened from the CHGV catalog [44] using CheckV [46] based a selection criterium of >90% completeness. In total, we selected 5,956 complete (with 100% completeness) and 2,892 high-quality (with >90% completeness) phage genomes for subsequent analyses.

### Identification and characterization of DNA methylation profiles of the 8,848 phages across 104 fecal samples

Methylated sites were identified for the 8,848 gut phage genomes across 104 fecal samples with SMRT sequencing data, by using the Base Modification Analysis Application module of the SMRT Link tool (v10.1) from https://www.pacb.com/support/software-downloads/ on Jan 15, 2022. For each sample, the consensus reads (HIFI reads) generated by the PacBio CCS mode were aligned to the 8,848 CHGV-HQ phage genomes and then the SMRTlink tool was used to detect the methylated bases which compared the mean Inter Pulse Duration ratio (IPDr) of all the subreads belonging to the aligned HIFI reads at a position of the reference genome with that of the unmethylated bases [15].

*MotifMaker* (SMRTLink, https://www.pacb.com/support/software-downloads/) was used to identify the methylated motifs, but did not product any results. HOMER [48] (v4.11), a popular motif prediction tool for analyzing ChIP-seq data, was also used to identify motifs; although it identified a lot of motifs from our data, HOMER did not report the exact locations on the viral genome. We speculate that these tools were designed for analyzing prokaryotic/eukaryotic genomes that are significantly longer than viruses, and thus might not be suitable for viral genome analysis. We thus identified methylation motifs using MEME-ChIP [25b] (v5.4.1; default parameters). Motifs were then dereplicated with a customized R pipeline (https://github.com/whchenlab/gutphagemethylome). To further remove false positives, we also required that the motifs should be found in at least 100 phage genomes. In total, 157 non-redundant motifs were identified (Table S1). A customized R pipeline was used to compare the motifs with those in the REBASE [26]. A rather relaxed criteria was used to search for the overlap; for example, motifs that are one of the possibilities of the motifs in public database are considered overlapped, e.g., AK<u>C</u>TCG is considered to be overlapped by B<u>C</u>NC (the former is one of the possibilities of the latter). In total, 155 were the same as compared with those in the REBASE (Table S1).

### Annotation of methylases (MTases) in phage genomes as well as bacterial/archaeal reference genomes (BRGs)

Prodigal [49] v2.6.3 with default parameters was used to predict genes from the 8,848 gut phage genomes. The predicted protein sequences were searched against the Conserved Domain Database (CDD) database [29] using RPS-BLAST (v2.12.0+; part of the BLAST package [21]; -evalue 1E-5). MTase proteins were identified with the following domain: cd21179.smp, COG0350.smp, KOG3191.smp, pfam05869.smp, pfam12047.smp, PRK10904.smp, PRK11524.smp, TIGR00589.smp, TIGR00675.smp, TIGR01712.smp and TIGR02987.smp. The MTases were further annotated by blast against the REBASE [26] with BLASTp (e-value <1e-5).

The same methods were used to identify MTase proteins in bacterial/archaeal reference genomes in the Unified Human Gastrointestinal Genome collection v2.0 (UHGG2) [23] and public phage genomes in the Gut Phage Database (GPD) [6b].

The search of MTases might limited by the CDD search method, for that it might underestimate/misclassify some of the MTase hits.

### Selection of high-quality phage genomes from GPD, the Gut Phage Database

For comparison purposes, 40,140 high-quality phage genomes (>90% completeness) were also selected from the Gut Phage Database (GPD) [6b] using CheckV [46]. Among which 17,743 (44.2%) were of 100% completeness; this subset of GPD genomes were referred as to GPD-HQ in this study.

### Machine learning models for prediction of human gut virome and performance evaluation

To check the quality and representativeness of CHGV-HQ and compare it with public human (gut) virome databases including GVD [6a], GPD [6b], MGV[6c], CHVD [6d] and DEVoC [6e], a series of neural network models were trained with the same architecture as DeepVirFinder [50].

For CHGV-HQ, GVD and DEVoC, we kept all sequences as the true positive datasets, while for GPD and MGV, the longest sequence of each VC (<u>V</u>iral <u>C</u>luster, considered as genus level of viruses) were selected as representative genomes considering memory consumption and training time, and referred to as GPD-rep and MGV-rep, respectively. As for CHVD, the intestine origin genomes were extracted. All above mentioned datasets were kept as positive datasets separately.

The bacterial genomes over 1.5kbp from human intestine were collected as negative samples from UHGG-Minus (Unified Human Gastrointestinal Genomes v2 [23] without prophage sequence; prophage identified with Phage_Finder [47] with default parameter).

We randomly selected 80% of each collection as training set, and other 20% as the test set. The DNA sequences in training set were consecutively segmented into non-overlapping fragments (1 kbs), then encoded into numerical matrices with a one-hot encoding method. The testing dataset were also segmented into non-overlapping fragments, and for each sequence, the average score of its fragments was assigned. For that the bacterial fragments were significantly out-numbered the viral ones, we randomly down-sampled the negative bacterial training set to match the positive viral one for every model.

We assessed the representativeness of CHGV-HQ with two experiments. The performance of the models we built can be considered as an indicator of the quality and representativeness of each dataset for that the same method was used.

In the first experiment, a test on an independent dataset, IMG/VR database [24] (https://img.jgi.doe.gov/cgi-bin/vr/main.cgi) and bacterial sequences from the test set of UHGG-Minus, were carried out. To make sure that the CHGV-HQ training set was independent with this IMG/VR test set, the pairwise comparison was conducted, by blasting (BLASTn[30], v2.2.26+; E-value < 1e-5) IMG/VR sequences against CHGV-HQ. The IMG/VR genome was kept only if it was with 90% nucleotide identity on less than 70% (calculated with bedtools v2.29.1 [51]) of its genome. The same procedure was carried out for the bacterial test dataset.

We then calculated metrics such as accuracy, precision, true negative rate(tnr), recall, f1-score, and AUC (Area Under the receiver operating characteristic Curve) score to evaluate the CHGV-HQ model.

In the second experiment, we applied the CHGV-HQ model to the above human gut virome datasets to see if most of the latter sequences could be correctly recognized.

In addition, viral-recognition models trained on the public datasets were also applied to the CHGV-HQ genomes, to further validate the latter’s viral identity.

### Estimation of the prevalence of the CHGV genomes at the viral contig

To estimate the abundance of phage genomes, the VLP-Next generation sequencing clean reads were mapped to the CHGV-HQ genomes using Bowtie2. Then, we calculated the reads per kilobase million (RPKM) value of each phage genome. An arbitrary abundance cutoff of 0.5 was used to define the presence of a phage in each of the 104 samples. Changing the presence/absence threshold up to 5 RPKM did not affect our main results.

### Partial correlation calculation between the phage prevalence and methylation densities with the sequencing depth control for

Partial correlation is a measure of the strength and direction of a linear relationship between two continuous variables whilst controlling for the effect of one or more other continuous variables. The partial correlations between phage prevalence and methylation densities, whilst controlling for sequencing depth were calculated using R (v 4.0.5) with “ppcor” Rpackage [52]. To plot the results after the sequence depth was controlled for, the residuals of the prevalence and methylation densities were calculated using the ‘resid’ function implemented in R with commands like ‘resid(lm(prevalence ∼ sequencing_depth))’ and ‘resid(lm(meth_density ∼ sequencing_depth))’.

### Annotation of crAssphages and Gubaphages

crAss-like phages were identified using the same method reported in a previous study (ref. [53]). First, phage nucleotide sequences were compared against the protein sequence of the polymerase (UGP_018) and the terminase (UGP_092) of the prototypical crAssphage (p-crAssphage, NC_024711.1) using BLASTx [21] (v2.2.26+). Second, the nucleotide sequence similarities between the phages and the p-crAssphage genome were assessed using BLASTn [21]. A phage was then classified as a crAssphage when it was longer than 70kb and met at least one of the following criteria:

1. had a BLASTx hit with an E-value <1e-10 against either p-crAssphage polymerase or terminase
2. showed ≥95% nucleotide identity over 80% of the contig length with the p-crAssphage genome

Gubaphages were identified by searching for the large terminase gene of the Gubaphage genomes obtained from the Gut Phage Database (GPD) [6b] using BLASTp (v2.2.26+, E-value < 1e-5). Gubaphage were then classified into four genera (G1.1, G1.2, G1.3 and G2) according to the similarity of the large terminase genes to those identified in the GPD.

We identified in total 245 crAssphage and 180 Gubaphage genomes in our dataset.

For comparison, 1,490 Gubaphages and 2,057 crAssphages were extracted according to the genome annotations in GPD-HQ.

### Annotation of phage lifestyles

Phage lifestyles were predicted using DeePhage [27] v1.0 with default parameters. DeePhage uses a scoring system to classify phage genomes into four categories, including temperate (with scores <=0.3), uncertain temperate (0.3∼0.5), uncertain virulent (0.5∼0.7), and virulent (>0.7). Higher scores indicate higher virulence [27]. Among the 8,848 phages, 1,211 (13.68%), 5,272 (59.58%), 1,647 (18.61%) and 718 (8.11%) were classified as temperate, uncertain temperate, uncertain virulent, and virulent respectively (Table S1).

### Clustering analysis of the phage- and bacteria-encoded MTase proteins

Protein sequences of MTases from phages, bacterial and archaeal genomes were merged and a BLASTp (v2.2.26+; E-value 1e-5) algorithm was used to search the merged dataset against itself for homologous sequences. The query-hit pairs were further filtered with a coverage >75% on the query proteins. The filtered BLASTp results were used as input for a Markov Clustering Algorithm [54] (MCL v14-137) with default parameters to generate protein clusters (PCs). The clustering analysis was performed separately for MTase proteins.

### Phage host prediction using MTases

To check if MTases can be used to correctly predict phage hosts, and determine the best parameters for the predictions, a published phage-host dataset was obtained from the Microbe-versus-Phage (MVP) database [3] and used as gold standard. Of note, only experimentally validated phage-host relationships and those inferred from prophages were retained for further analysis, resulting in 778 relationships between 422 phages and 1517 prokaryotic hosts (Table S1). MTase proteins were identified for the phages and prokaryotic hosts, as described in the previous section. BLASTp (v2.2.26+) [21] was used to detect homologous relationships between phage- and bacteria-encoded MTases. We calculated a sequence similarity score (BLASTp q-identity × query coverage, SimScore) and used AUC scores to evaluate accuracy of the SimScores to distinguish true interactions from randomly generated non-interacting phage-host pairs. Such an analysis can help us to determine the best SimScore cutoffs by taking specificity, sensitivity or both into consideration. To predict hosts for the phages, phage-encoded MTase protein sequences were searched against those of the UHGG2 using BLASTp (v2.2.26+). A bacterial/archaeal genome whose BLASTp SimScore to a phage MTase protein was higher than the threshold was assigned as the phage host. To avoid wrong host assignments because of the lack of bacterial reference genomes, we adopted a rather stringent threshold to assign a bacterial host to a phage if i) at least two phage-encoded MTase proteins had SimScore higher than 90% with the host-encoded counterparts, and ii) at least two such sequence-matches agreed on the same host. After filtration, the annotation for each protein with highest BLASTp SimScore were kept. At these criteria, we achieved precision of 73.5% and 85.2% at the species and genus levels respectively on the MVP dataset (Fig. 4D). The predicted phage-host relationships can be found in Table S1.

### Host range calculation of host-prediction results

Host prediction results were evaluated by calculating the host range for each phage. For phages with only one predicted host, the host range was at the species level. For those with multiple predicted hosts, the host range was calculated as the last common ancestor (LCA) of all the hosts on the NCBI taxonomic database using an in-house R script (https://github.com/whchenlab/gutphagemethylome.).

### Statistics and other bioinformatics analyses

All processed data, if not otherwise stated, were loaded into R (v4.0.5, https://www.r-project.org/), analyzed or visualized.

## Availability of data and materials

The raw sequencing data used in this study are available in the CNCB GSA database under accession code PRJCA008836 (accessible via either the GSA link https://ngdc.cncb.ac.cn/gsa/browse/CRA006494 or the BioProject page https://ngdc.cncb.ac.cn/bioproject/browse/PRJCA008836).

The software parameters and scripts, genomic data used in the analysis process have been uploaded to a Github repository at https://github.com/whchenlab/gutphagemethylome.

## Acknowledgements

We thank all members of the Chen, Liu and Zhao labs for their help with the fecal sample collection, and insightful discussions.

## Ethics approval and consent to participate

This study was approved by the Ethics Committee of the Tongji Medical College of Huazhong University of Science and Technology (No, S1241) and the Human Ethics Committee of the School of Life Sciences of Fudan University (No, BE1940).

## Funding

This research is supported by National Natural Science Foundation of China (32070660 to W.H.C; 61932008, 61772368 to X.M.Z; 31770132, 81873969 to Z. L), National Key Research and Development Program of China (2020YFA0712403 to X.M.Z; 2019YFA0905600 to W.H.C and Z. L), NNSF-VR Sino-Swedish Joint Research Programme (82161138017 to W.H.C), and Shanghai Municipal Science and Technology Major Project (2018SHZDZX01 to X.M.Z).

## Author Contributions

WHC, XMZ, ZL and PB designed and directed the research; CS did most of the analysis; JC managed the sampling and did some of the experiments; XZ, MJ, and YL helped with the experiments; MJ also helped with the sample collection and phage enrichment experiments; YD did the machine learning analysis. CS wrote the paper with results from all authors; WHC, XMZ, ZL and PB polished the manuscript through multiple iterations of discussions with all authors. All authors have read and approved the final manuscript.

## Competing interests

The authors declare no competing interests.

## Notes

https://ngdc.cncb.ac.cn/bioproject/browse/PRJCA008836

### Competing Interest Statement

The authors have declared no competing interest.

