## Supplementary figures and info for "Long-read sequencing reveals extensive DNA methylations in human gut phagenome contributed by prevalently phage-encoded methyltransferases"

### Supplementary Tables

Table S1. A list of 9401 phages and related information in details.
Table S2. MTases identified from the 9401 phages, UHGG2 genomes, and their clustering results

### Supplementary Figures

#### Figure S1

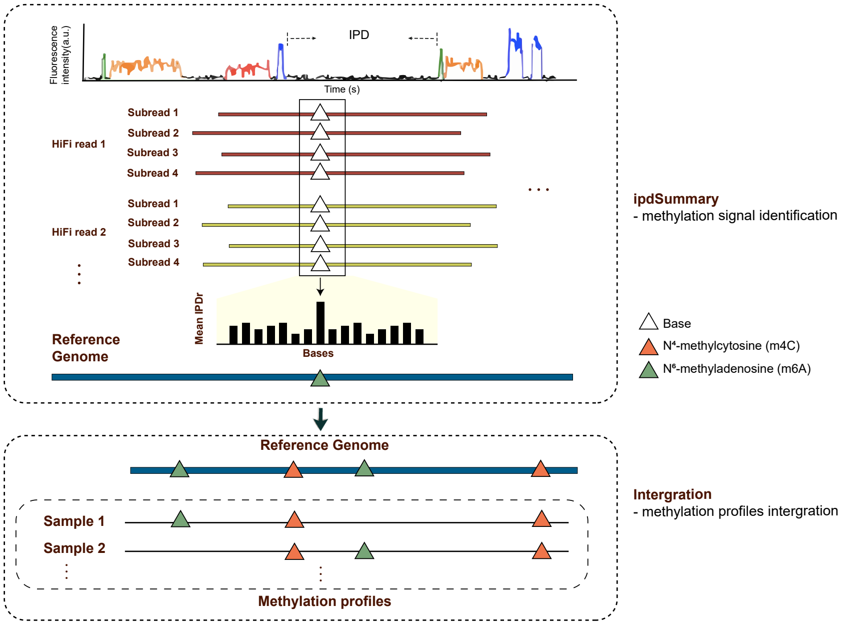


**Figure S1,** DNA methylation identification using SMRT sequencing.

### Figure S2

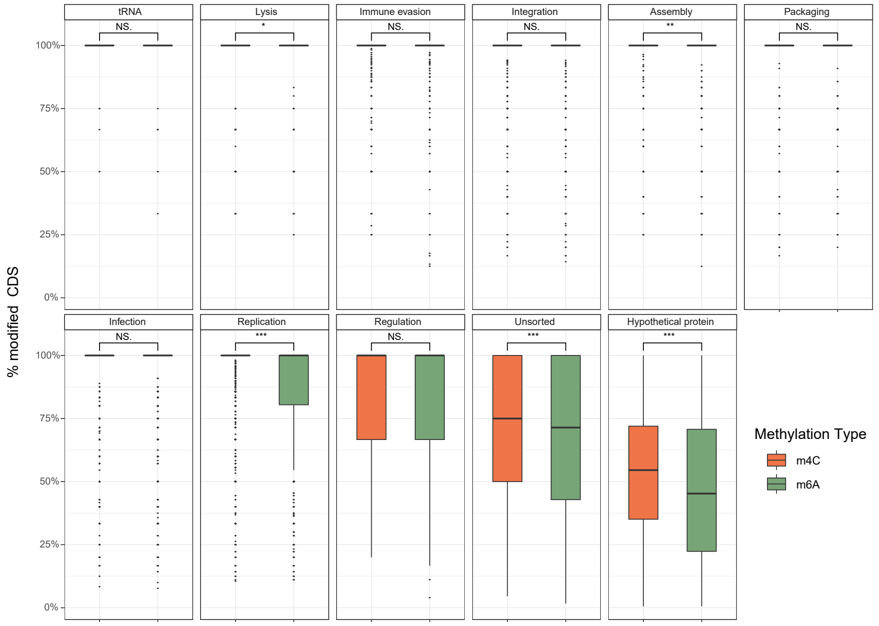


**Figure S2,** Differential distribution patterns of m6A and m4C modifications in coding genes with different functions.

### Figure S3

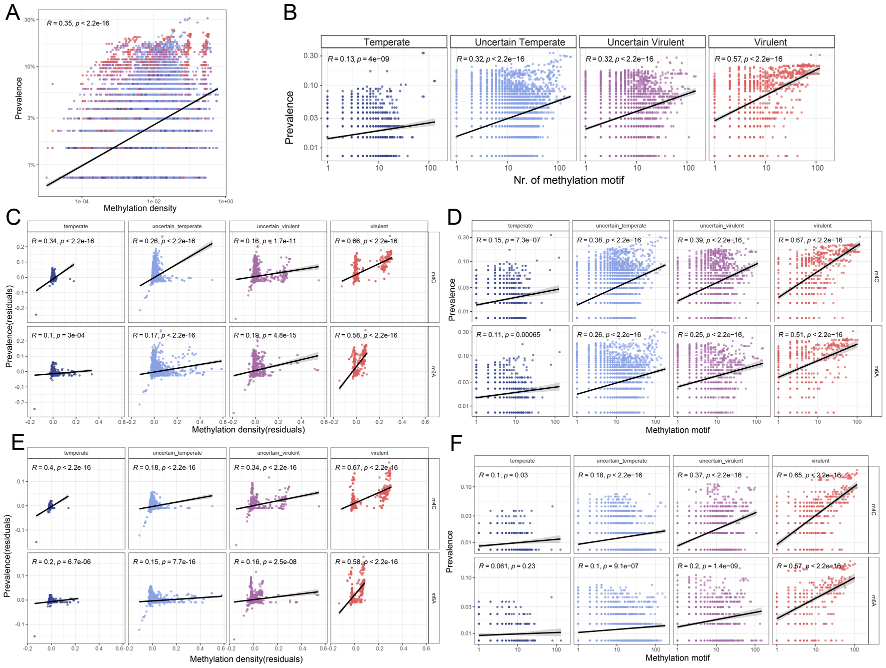

**Figure S3,** The fitness of the CHGV-HQ phages, measured by the prevalence (lower panels) across 104 fecal samples, was positively correlated with **A)** overall DNA methylation density, **B)** the total numbers of methylation motifs, **C)** overall DNA methylation density of m4C and m6A and **D)** numbers of methylation motifs of m4C and m6A. **E,F)** The prevalence was calculated by using an abundance cutoff of 5 as the presence/absence threshold, showing that changing the abundance cutoff did not affect our main results

### Figure S4

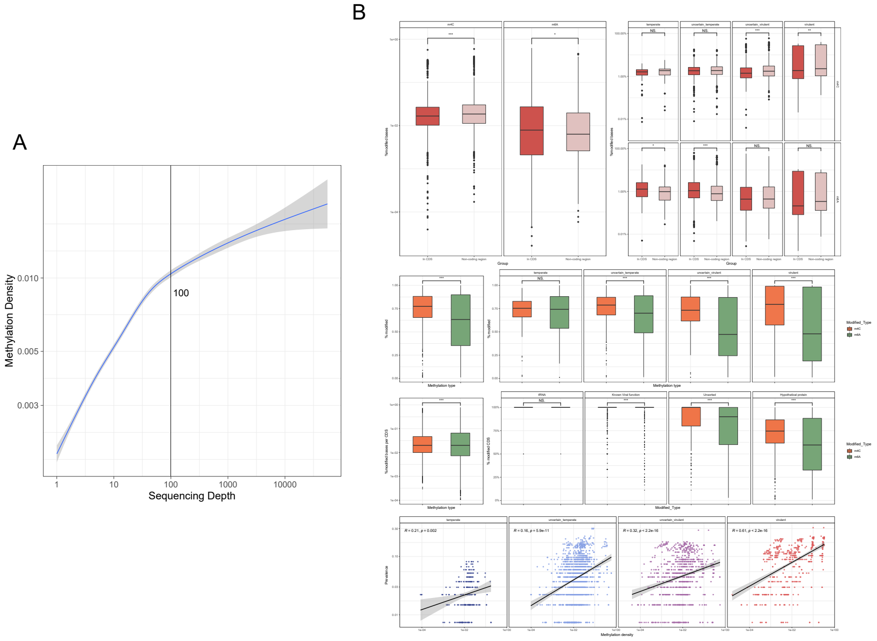


**Figure S4, A)** The rarefaction of phage methylation density with increasing TGS coverage. The lines represent the coverage under 100. The methylation density among CDS and non-CDS region, methylation density among different gene functions, and phage fitness under the coverage cutoff of **B)** 100. Limiting our analysis to phages with long-reads coverages did not affect our result.

#### Figure S5

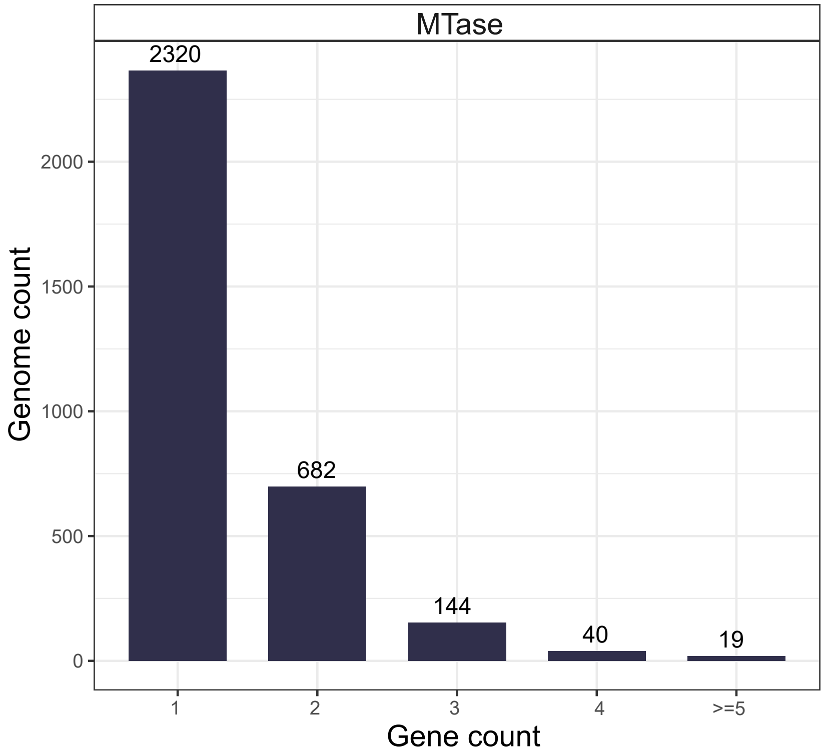


**Figure S5,** The number of genomes encode different count of MTases. Most phages contain one MTase gene, but some can encode multiple ones.

### Figure S6

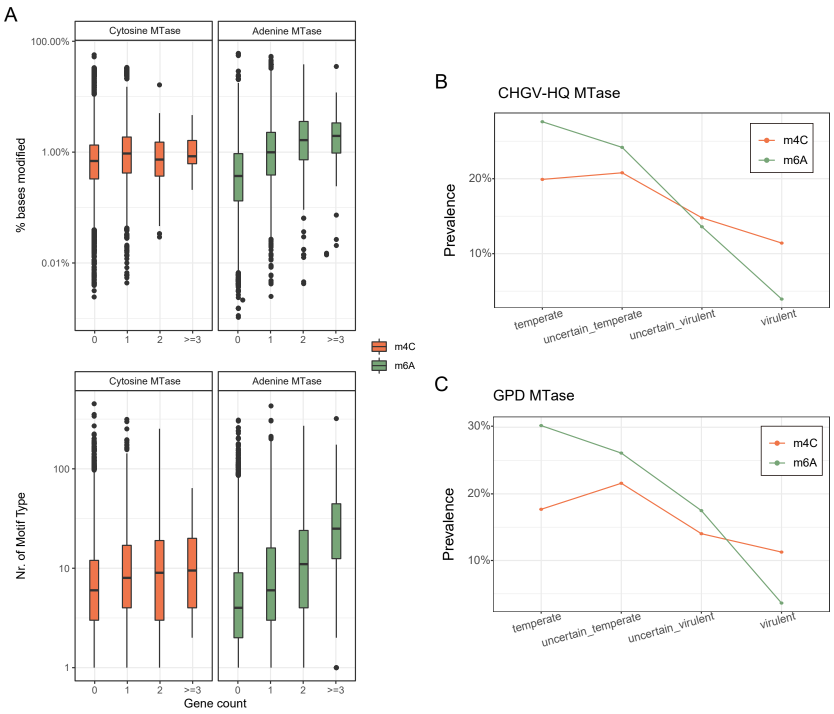


**Figure S6, A)** Both the methylation densities and the numbers of methylation motifs increased with increasing number of phage-encoded MTases responsible for individual modification types. A higher prevalence of MTase genes from CHGV **B)** and GPD **C)** is associated with decreasing phage virulence. The trends in the individual MTase types, i.e., MTases responsible for m4C and m6A modifications were largely the same.

### Figure S7

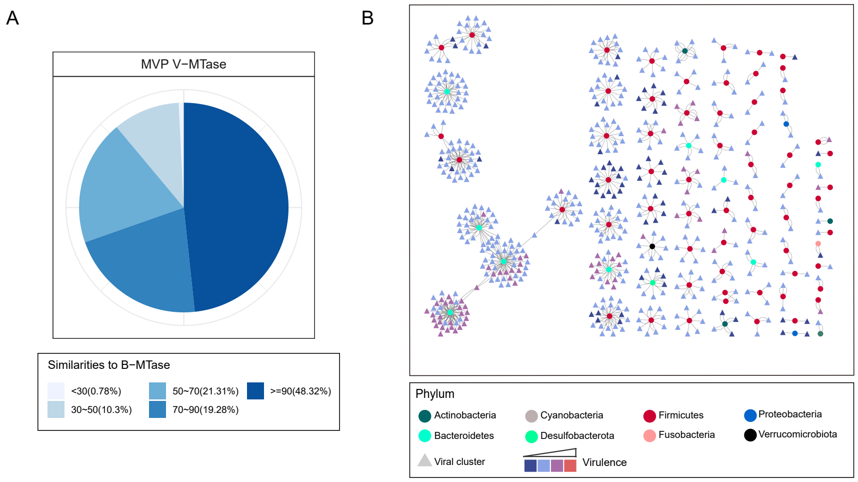


**Figure S7, A)** 48.32% of the MVP V-MTases share over 90

### Figure S8

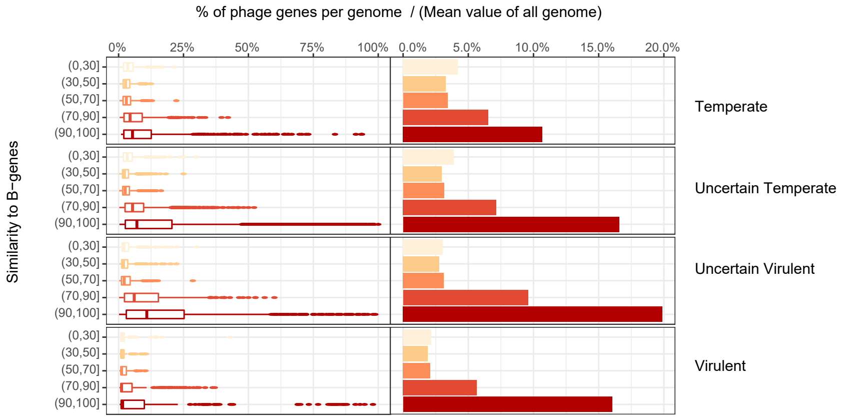


**Figure S8** 20% genes per phage genome (V-genes) share significant protein similarities with the UHGG2 gut bacteria(B-genes). The trends stay the same among different lifestyles.

### Figure S9

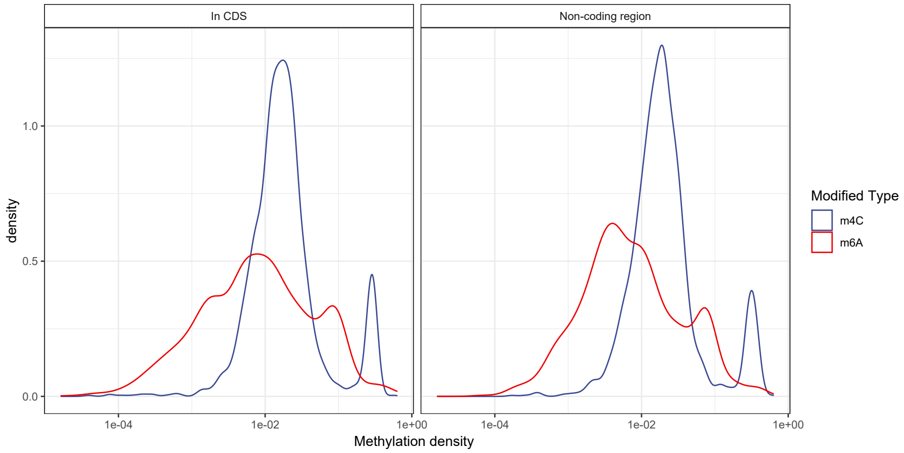


**Figure S9** Most methylation-positive genomes are with high methylation density, no matter coding and non-coding regions.
